# Benzoquinones in the defensive secretion of a bug (*Pamillia behrensii*): a common chemical trait retrieved in the Heteroptera

**DOI:** 10.1101/2020.12.11.421891

**Authors:** Julian M. Wagner, Thomas H. Naragon, Adrian Brückner

## Abstract

Benzoquinones are a phylogenetically widespread compound class within arthropods, appearing in harvestman, millipedes and insects. Whereas the function of benzoquinones as defensive compounds against potential predators and microbes has been well established, the full extent of benzoquinone usage across arthropods, and especially within Insecta, has yet to be established. Adding to the growing list of unique evolutionary origins of benzoquinone employment, we describe in this paper the metathoracic scent gland secretion of the mirid bug *Pamillia behrensii*, which is composed of heptan-2-one, 2-heptyl acetate, 2,3-dimethyl-1-4-benzoquinone, 2,3-dimethyl-1-4-hydroquinone as well as one unknown compound. Similarly, to many other arthropods that use benzoquinones, *Pamillia* releases the contents of its gland as a defensive mechanism in response to harassment by other arthropod predators. Morphological investigation of the gland showed that the benzoquinone-producing gland complex of *P. behrensii* follows a similar blueprint to metathoracic scent glands described in other Heteropterans. Overall, our data further underpins the widespread convergent evolution and use of benzoquinones for defense across the Arthropoda, now including the order Hemiptera.

## Introduction

Chemical convergence is the evolution and widespread use of certain compound classes by different taxa for similar purposes (Beran et al. 2019; Brückner and Parker 2020). Evolutionary patterns of this nature suggest potential genetic biases at the biosynthetic level or the efficiency of certain compound classes at eliciting a given response in other organisms within certain ecological settings as drivers of selection (Brückner and Parker 2020; Chevrette et al. 2020). ‘

One striking example of chemical convergence is the frequent evolution of aromatic benzoquinones (BQs) as defensive compounds in arthropods (**Fig 1A**). The past 60 years of research has uncovered a smorgasbord of quinone containing glands in insects, harvestment, and millipedes (e.g., Blum 1981; Meinwald et al. 1966; Raspotnig et al. 2017; Roth and Stay 1958; Shear 2015). In insects, BQs have a broad taxonomic distribution, spanning both hemi- and holometabolous insects, with representatives in the earwigs (Dermaptera), lubber grasshoppers (Orthoptera), cockroaches (Blattodea), caddisflies (Trichoptera) and most prominently beetles (Coleoptera) (Blum 1981; Eisner et al. 2005; Francke and Dettner 2005).

**Figure 1.**
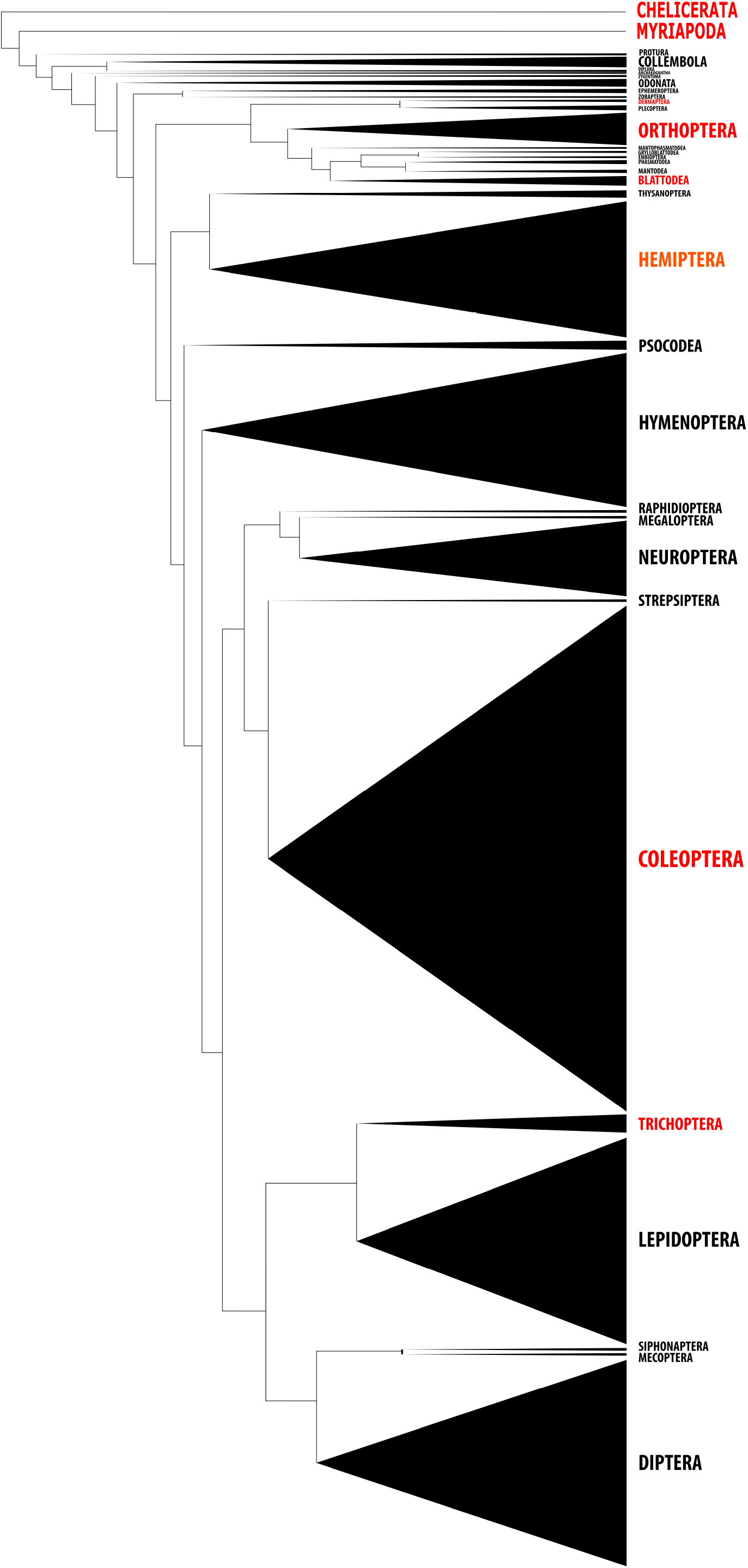
Convergent evolution of benzoquinones across the arthropods. Conceptual phylogenetic tree of arthropods - focusing on insects - depicting taxa producing benzoquinones based on previous research (in red) and results presented in this study (in orange). The tree topology is based on Misof et al. (2014) and the group size is proportional to the number of species based on Stork (2018).

Despite the similarity of the chemical phenotype, it is yet to be determined if the biosynthetic pathways for the BQs are also convergent (Blum 1981; Morgan 2010). Some have argued that BQs may be produced by enzymes related to tyrosine dependent cuticle tanning (Duffey 1974; He et al. 2018; Roth and Stay 1958), while others have demonstrated the possibility of *de novo* BQ synthesis from poly-β-carbonyls via head-to-tail condensation (Meinwald et al. 1966; Morgan 2010; Rocha et al. 2013). In contrast, the function of BQs in chemical defense against predators and as antimicrobial agents is non-controversial and widely documented (e.g., Eisner et al. 2005; Gasch et al. 2013; Li et al. 2013; Ruther et al. 2001; Shear 2015).

Here we provide another example for the widespread evolution and chemical convergence of benzoquinones, and report on the first evidence of this compound class in the Heteroptera (Aldrich 1988). We chemically analyzed metathoracic scent gland (MTG) secretion of the mirid bug *Pamillia behrensii*, Uhler (**Fig 2D**), investigated the internal and external morphology of the gland via confocal microscopy and investigated the biological function of MTG compounds.

**Figure 2.**
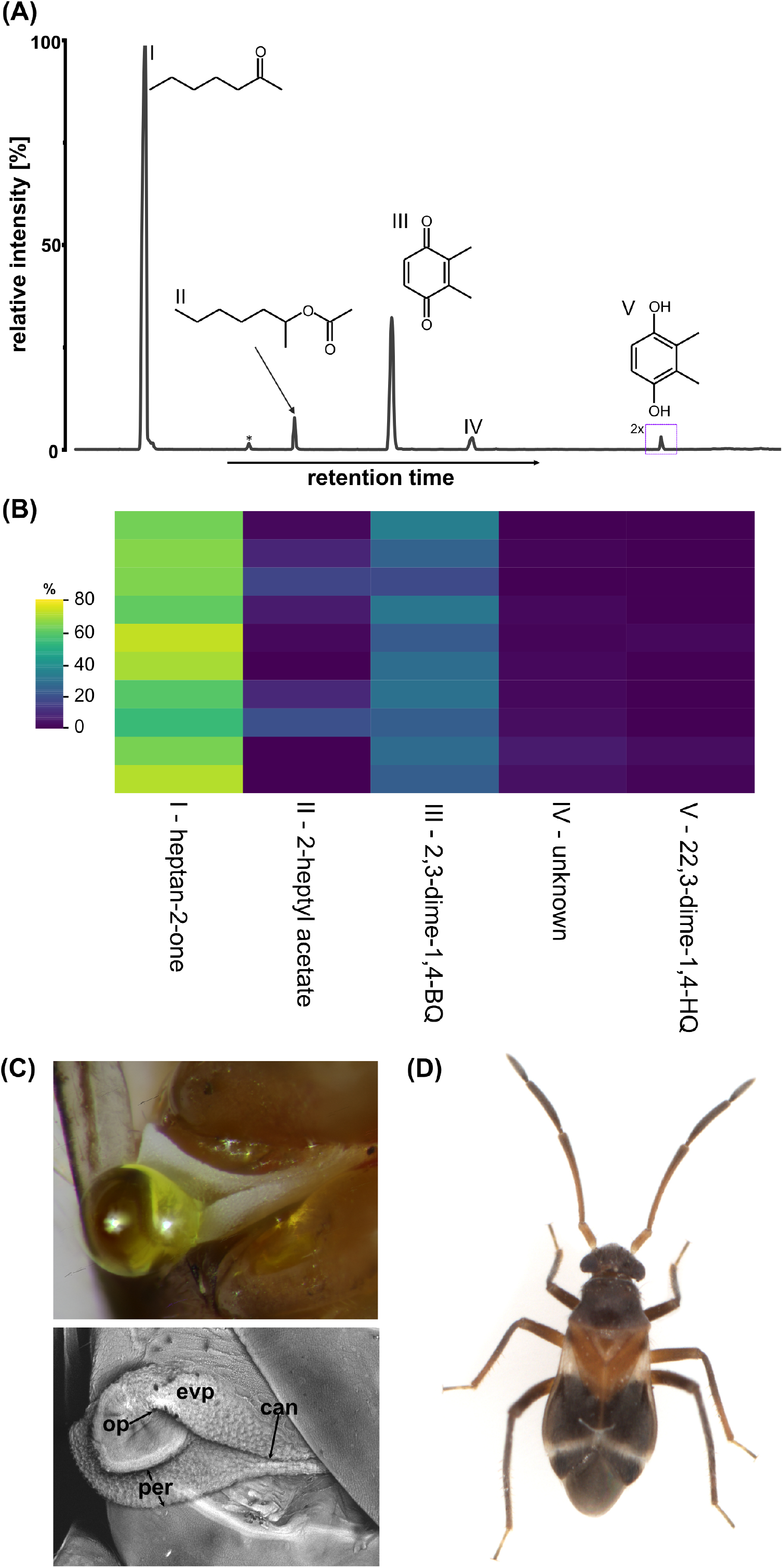
Metathoracic scent gland (MTG) chemistry of *Pamillia behrensii*. **A:** Example GC traces of *P. behrensii* gland exudates. Compounds are labeled in the trace and mass spectrometric data is detailed in Table 1. **B:** Matrix plot shows the composition of eleven individuals. The gland compounds in the order of their retention indices are heptan-2-one, 2-heptyl acetate, 2,3-dimethyl-1,4-benzoquinone, unknown IV and 2,3-dimethyl-1,4-hydroquinone. The legend indicates the percentage composition for each compound per specimen. **C:** Outer morphology of MTG of *P. behrensii*. The top image shows a droplet of the glandular secretion shortly after gland use. The yellow color indicates the presence of benzoquinones. The bottom image details the surface morphology around the gland opening. op= gland opening; evp= evaporation surface; per= peritreme; can= canal from ostiole to coxal pit. **D:** Habitus image of *P. behrensii* with scale bar.

## Materials and Methods

### Collection

Individual *Pamillia behrensii* were collected in close proximity to nests of the velvety tree ant *(Liometopum occidentale*) on October 23rd 2019 and November 17th 2020 at Chaney Canyon, Altadena, CA (34°13’01.7”N 118°09’15.1”W). Insects were carefully transferred to the laboratory in tubes lined with moist tissue paper and directly used for chemical analysis or imaging.

### Chemical analysis

To identify compounds stored in the metathoracic scent gland (MTG) of *Pamillia behrensii* we used whole body extraction in 50 μl hexane for 10 min, which is a well-established method in Heteropterans (e.g., Leal et al. 1994; Zarbin et al. 2000). Crude hexane sample aliquots (1 μl) were analyzed on a GCMS-QP2020 gas chromatography/mass-spectrometry system (Shimadzu, Kyōto, Japan) equipped with a ZB-5MS fused silica capillary column (30 m x 0.25 mm ID, df= 0.25 μm) from Phenomenex (Torrance, CA, USA). Samples were injected using an AOC-20i autosampler system from Shimadzu into a split/splitless-injector operating in splitless-mode at a temperature of 310°C. Helium was used as the carrier-gas with a constant flow rate of 2.13 ml/min. The chromatographic conditions were as follows: The column temperature at the start was 40°C with a 1-minute hold after which the temperature was initially increased 30°C/min to 250°C and further increased 50°C/min to a final temperature of 320°C and held for 5 minutes. Electron impact ionization spectra were recorded at 70 eV ion source voltage, with a scan rate of 0.2 scans/sec from *m/z* 40 to 450. The ion source of the mass spectrometer and the transfer line were kept at 230°C and 320°C, respectively.

We integrated chromatograms manually using LabSolutions Postrun Analysis (Shimadzu, Kyōto, Japan), quantified the ion abundance and calculated the relative composition of individual compounds compared to the total ion abundance. Compounds were identified based on the *m/z* fragmentation patterns and retention indices (RIs) calculated after Van den Dool and Kratz (1963) using a standard alkane mixture. Synthetic heptan-2-one was purchased from Sigma-Aldrich (St. Louis, MO, USA) and natural 2-ethyl-1-4-benzoquinone and 2-ethyl-1-4-hydroquinone from the secretion of *Tribolium castaneum* (Li et al. 2013) were used as reference compounds. 2-Heptyl acetate was synthesized by mixing acetic acid with an excess of 2-heptanol and H_2_SO_4_ (all Sigma-Aldrich) as catalyst and subsequently heating the mixture for 48 h at 45 °C. The crude ester was extracted with hexane and the organic layer was washed with water, saturated Na_2_CO_3_ solution (2x) and saturated NaCl solution and finally dried over anhydrous Na_2_SO_4_.

### Histochemistry and imaging

Adult bugs were immersed in PBS and legs were removed with forceps. The dorsal parts of the thorax were removed by cutting around the abdominal margin with dissection scissors and the ventral thorax was fixed in 4% paraformaldehyde (25 minutes, room temperature), washed in PBS+0.02% Triton X100, and subsequently stained with Alexa-647-Phalloidin (Thermo Fisher, Waltham, MA, USA) to label muscles and Alexa-546-WGA (Thermo Fisher, Waltham, MA, USA) to stain membranes. Metathoracic scent glands from both sides were imaged as whole mounts of ventral thoraxes in ProLong Gold Antifade Mountant (Thermo Fisher, Waltham, MA, USA), using a Zeiss LSM 880 (Carl Zeiss, Jena, Germany) with airyscan.

### Behavioral assay

To confirm that P. behrensii deploys gland contents in response to threat we performed a behavioral trial in tandem with solid phase microextraction (SPME). A 4-cm circular behavioral arena was constructed from 1/8th inch infrared transmitting acrylic (Plexiglass IR acrylic 3143) with a port on the side for a 65 µm polydimethylsiloxane/divinylbenzene fiber from Supelco (Sigma-Aldrich; St. Louis, MO) to sample released volatiles. For compound desorption the fiber was placed in the injector port for 1 min operating at 230°C and the GC/MS run was carried out as outlined below. After each trial the fiber was baked for 30 min at 230°C. SPME-GC/MS runs were used to identify the fraction of volatile metathoracic scent gland (MTG) compounds (see e.g. Krajicek et al. 2016).

Behavior was recorded for thirty minutes at 25 fps with a machine vision camera (Flir BFS-U3-16S2M-CS). Bugs were anesthetized on ice and placed in an arena either alone, with five *L. occidentale*, or with five *L. occidentale* with superglued mandibles to reduce ant aggressiveness. To assess behavior, bugs were labeled with DeepLabCut (DLC) to quantify movement over time (Mathis et al. 2018). Median filtered, labeled bug locations were used to sum to total distance traversed during the trial. SPME-GC/MS runs were used to identify the fraction of volatile metathoracic scent gland (MTG) compounds (see e.g. Krajicek et al. 2016). A linear regression was performed on 10,000 bootstrap samples of the data in python to calculate confidence intervals for the regression slope. A Jupyter notebook is available which reproduces the analysis at https://github.com/julianmwagner/bq_bug.

## Results and Discussion

In total, we found five compounds in the MTG extracts of *Pamillia behrensii* of which we were able to identify four (**Fig 2A; Tab 1**). Compound I elicited a prominent fragment ion at *m/z*= 58 arising from McLafferty rearrangement, which is characteristic for saturated methyl-ketones, together with a molecular ion at *m/z*= 114 we identified I as heptan-2-one and eventually confirmed our identification with a synthetic standard.

**Table 1.** Gas chromatographic and mass spectrometric data of the metathoracic scent gland secretion released by the mirid bug *Pamillia behrensii*. Molecular ions (M+) are marked in bold. Retention indices (RI) were calculated according to van den Dool and Kratz (1963).

| RI | RI literature | Identified as | Mass spectrometric fragmentation *m/z* (relative intensity) | Secretion profile [%] |
| --- | --- | --- | --- | --- |
| 890 | 892<sup>a</sup> | heptan-2-one | 114 (M<sup>+</sup>, 4), 99 (3), 85 (3), 71 (15), 58 (55), 43 (100) | 64.1±5.2 |
| 1041 | 1043<sup>b</sup> | 2-heptyl acetate | 158 (M<sup>+</sup>, 1), 115 (3), 102 (2), 98 (7), 87 (22), 70 (8), 69 (7), 61 (3), 56 (15), 43 (100) | 6.5±6.5 |
| 1118 | 1119<sup>c</sup> | 2,3-dimethyl-1-4-benzoquinone | 136 (M<sup>+</sup>, 85), 108 (71), 107 (59), 82 (59), 80 (31), 79 (76), 54 (100), 53 (44) | 26.5±4.5 |
| 1223 | - | unknown | 138 (M<sup>+</sup>, 52), 110 (14), 95 (16), 82 (29), 67 (40), 54 (100) | 2.1±1.6 |
| 1432 | 1433<sup>c</sup> | 2,3-dimethyl-1-4-hydroquinone | 138 (M<sup>+</sup>, 100), 137 (25), 123 (53), 95 (29), 91 (25) | 0.8±0.7 |

<sup>a</sup>RI as reported by Schaidler et al. (2018)
<sup>b</sup>RI as reported by Bertoli et al. (2011)
<sup>c</sup>RI as reported by Rocha et al. (2013)

Compound II showed a molecular ion at *m/z*= 158 and a base ion at *m/z*= 43 indicating an acetate ester [additional R’COOH_2_ at *m/z*= 61; Urbanová et al. (2012)] with a heptanol alcohol moiety (ion at m/z= 115). Both diagnostic ions at *m/z*=87 and the pair at *m/z*=69/70 indicate a possible heptan-2-ol moiety (**Tab 1**). Hence, we tentatively identified II as 2-heptyl acetate (1-methylhexyl acetate) and eventually confirmed the compound’s identity by synthesis.

Compound III showed molecular ions at *m/z=* 136 as well as diagnostic ions at *m/z*= 108, *m/z*= 82, *m/z*= 79 and *m/z*= 54, indicating either a 2,3-dimethyl- or 2-ethyl-substituted 1-4-benzoquinone. An ion at *m/z*= 107 (i.e. 5-methyl-4-methylidenecyclopentenone ion) and III’s retention index (**Tab 1**) were, however, more indicative of 2,3-dimethyl-1-4-benzoquinone. Compound V showed a molecular ions at *m/z=* 138 (also the base ion) and diagnostic ions at *m/z*= 137 *m/z*= 95, as well as *m/z*= 91 which together with V’s retention index (**Tab 1**), were all indicative of 2,3-dimethyl-1-4-hydroquinone. We used the secretion from defensive stink gland of *Tribolium castaneum* (Li et al. 2013) which contains 2-ethyl-1,4-benzoquinone and 2-ethyl-1,4-hydroquinone as a natural standard and found no correspondence to III and V. Hence, we eventually identified III as 2,3-dimethyl-1,4-benzoquinone and V as 2,3-dimethyl-1,4-hydroquinone.

For compound IV we did not find any close hit in any compound library used (NIST14, Wiley-NIST 2009, FCSN2) and the identity of this compound remains to be elucidated (see **Tab 1** for *m/z* fragmentation). The MTG extracts of *P. behrensii* (**Fig 2A**), overall showed a very consistent chemical composition, with heptan-2-one always appearing in the greatest abundance, 2,3-dimethyl-1,4-benzoquinone at an intermediate abundance and 2-heptyl acetate, unknown IV as well 2,3-dimethyl-1-4-hydroquinone being minor compounds (**Fig 2B; Tab 1**).

Based on synthetic heptan-2-one and 2-methyl-1,4-benzoquinone, as external standards, we estimated 14.1±5.8 µg of heptan-2-one and 4.9±2.2 µg of 2,3-dimethyl-1,4-benzoquinone per individual, respectively. Based on the composition and amounts, we interpreted heptan-2-one as a solvent for the more noxious benzoquinone and hydroquinone, while the ester 2-heptyl acetate may be an additional repellent or act as surfactant as previously described for glandular secretions of rove beetles (Dettner 1984; Steidle and Dettner 1993).

Detecting 2,3-dimethyl-1,4-benzoquinone in the MTG extracts of a heteropteran (**Fig 2A**), adds another order of arthropods to the list of BQ-producers and underpins the remarkable pattern of chemical convergent evolution of BQs. Even though other aromatics compounds (e.g. vanillin, 2-phenylethanol, benzyl alcohol, or p-hydroxbenzaldehyde) have been described in other bugs, benzoquinones add another compound class to the chemical skillset of the Heteroptera. Focusing on aliphatic compounds, MTG secretions of bugs are typically composed of n-alkanes, alkenals, alkenyl alkanoates and occasionally monoterpenoids. While heptan-2-one appears to be a rather unusual compound for bugs as well, methyl ketones are common in arthropods. For instance, in ants, other hymenopterans, trichopterans as well as cyphophthalmid and dyspnoan harvestman they serve as alarm and sex pheromones or as defensive compounds (Blum 1981; Cheng et al. 2017; Löfstedt et al. 2008; Raspotnig et al. 2005; Schaider et al. 2018).

2,3-Dimethyl-1-4-hydroquinone in the MTG secretions of *P. behrensii* (**Fig 2A**) can be interpreted as a precursor molecule that is oxidized to the final benzoquinone *via* a laccase enzymes, as a recent transcriptomic study in termite soldiers suggested (He et al. 2018). Whether the hydroquinone is derived from tyrosine or was produced utilizing a polyketide-like mechanism (Brückner and Parker 2020) is, however, unknown.

As is typical for chemical release from the metathoracic scent gland of other bugs, we found that the yellow, quinone secretion is also expelled from this gland complex as small droplets (**Fig 2C**). The MTG was located between thorax segment II and III (**supplement Fig S1A**) and generally showed an outer morphology comparable to other Heteropteran glands (Aldrich 1988; Gonzaga-Segura et al. 2013; Hepburn and Yonke 1971): two adjacent cuticular folds form flaps/lips on the anterior and posterior edges of the gland opening (**Fig 2C**). The edges of the cuticular folds constitute the ostiolar peritreme, which likely aids in carrying the glandular secretions away from the opening and fosters increased compound dispersion (**Fig 2C**). Additionally, the roughly sculpted cuticle around the gland opening serves as an evaporation surface (**Fig 2C**), increasing the volatility of expelled MTG compounds (Aldrich 1988; Hepburn and Yonke 1971).

The internal part of the MTG complex **(Fig S1B and C**) consists of a lateral extension which connects the gland opening to the median reservoir where the secretions produced by primary gland are stored (Aldrich 1988; Hepburn and Yonke 1971). Despite the capability of the MTG of *P. behrensii* to synthesize quite unusual Heteropteran compounds, the general outer and inner morphology of the MTG complex appears to be similar to that found in other MTG containing species and we did not find any specialized adaptions to house these compounds.

To confirm *P. behrensii* uses its MTG secretion (**Fig 2A**) as a defensive compound, we performed behavioral experiments coupled with chemical headspace profiling with solid phase microextraction. We expected the chemical secretion expelled from the gland should function as an allomone in chemical defense, as quinones are known to be noxious and serve as predator repellents in many other arthropods (e.g., Eisner et al. 2005; Gasch et al. 2013; Li et al. 2013; Ruther et al. 2001). Like other chemicals, benzoquinones activate the transient receptor potential ankyrin 1 (TRPA1), a nonselective cation channel, which has a conserved function as a noxious chemical receptor in animals (Arenas et al. 2017; Ibarra and Blair 2013). TRPA1 might thus be a major target of many convergently evolved defensive chemicals like benzoquinones and initiate protective behavior after a chemical attack (Blair et al. 2016; Ibarra and Blair 2013).

We found direct evidence of active usage of the MTG gland for defense in our behavioral recordings. In cases where ants successfully grabbed the bug, deployed gland content could be seen smeared on the behavioral arena (**Fig 3A**). The compounds quickly evaporated off the arena surface and were readily measurable *via* SPME-GCMS. We additionally profiled the correlation of amount of benzoquinone deployed with bug movement in the behavioral arena (**Fig 3B**). We found a positive correlation (95% confidence interval of slope: 0.154-1.75) of BQ amount with the distance the bug traveled during the behavioral trial (a proxy bug agitation/escape response). This behavioral readout supports the idea that *P. behrensii* deploys its gland to confuse or repel its predators before fleeing. We thus demonstrated that *P. behrensii* uses the compounds stored in its MTG as a repellent against predators, which again underpins the chemical convergence of benzoquinones.

**Figure 3.**
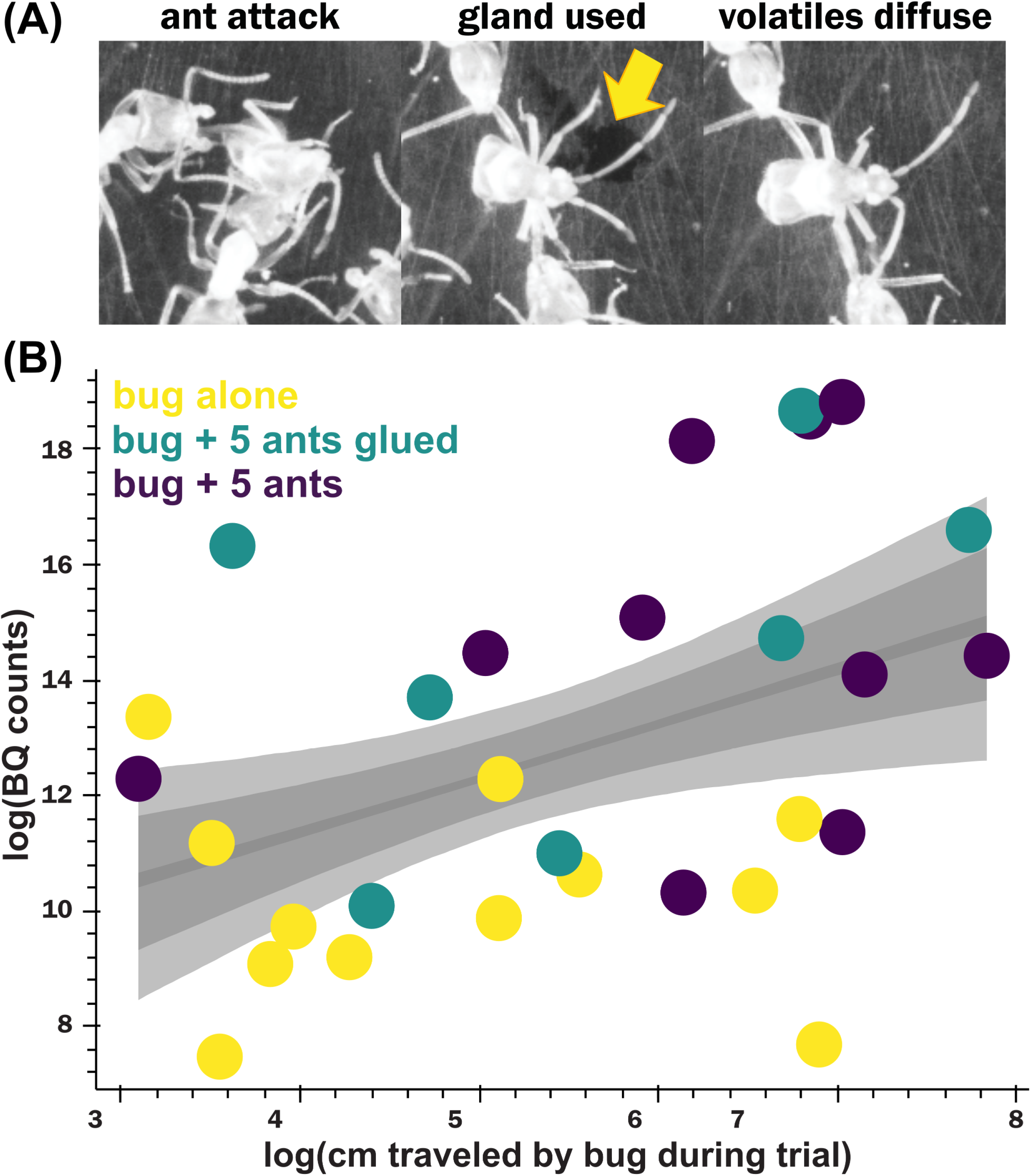
Behavioral evidence active defense gland use against predators. **A:** When placed in a behavioral arena ants attack *Pamillia behrensii* (left panel of A). In response to aggression, *P. behrensii* deploys defensive gland secretion, which can be seen by eye (A central panel, indicated with arrow). The gland secretion volatilizes, diffusing to nearby attackers (A right panel); volatile compound identity was confirmed *via* SPME-GCMS. **B:** Quantitative analysis of behavior indicated a significant positive correlation of bug movement (a proxy for bug agitation/attempts to flee predators) with benzoquinone quantity deployed. Grey intervals represent 95th, 75th and 10th percentile for regression confidence intervals from 10,000 bootstrap replicates. Together, these suggest that *P. behrensii* employs a joint strategy of gland deployment with an escape response to evade aggressive predators.

We found first evidence of a benzoquinone in the Heteroptera (Aldrich 1988; Morgan 2010). More specifically we detected 2,3-dimethyl-1-4-benzoquinone produced by and stored in the metathoracic scent gland complex of the mirid bug *Pamillia behrensii* and showed that it uses this BQ in combination with three other compounds for chemical defense against predators. Our study therefore highlights the remarkable convergent evolution of benzoquinones as defensive compounds across the Arthropoda, now including true bugs.

## Supporting information

Metadata Table

## Acknowledgment

We thank Joe Parker for making his laboratory available to us. JW was supported by NN. THN received a NN. AB is Simons Fellow of the Life Sciences Research Foundation (LSRF).

## Ethics statement

There are no legal restrictions on working with bugs. Field collection permissions were issues by California Department of Fish and Wildlife and the Angeles National Forest (US Forest Service; USDA).

## Authors contributions

JMW, THN and AB design research; JMW and THN collected specimens; THN performed microscopy; JMW performed and analyzed behavioral assays; JMW collected chemical data; AB analyzed chemical data and performed organic synthesis; AB supervised the project; AB wrote the paper with input from THN and JMW. JMW and THN contributed equally. All authors gave final approval for publication.

---

**Supplementary Figure S1** Overview of the morphology of the *P. behrensii* gland. **A:** Location of the metathoracic scent gland (MTG) opening located between thorax segment II and II. **B** + **C:** Inner morphology of the MTG showing the median reservoir where the secretions produced by primary gland are stored.

