## Supplementary figures and images for "Benzoquinones in the defensive secretion of a bug (*Pamillia behrensii*): a common chemical trait retrieved in the Heteroptera"

### Supplemental Figure S1

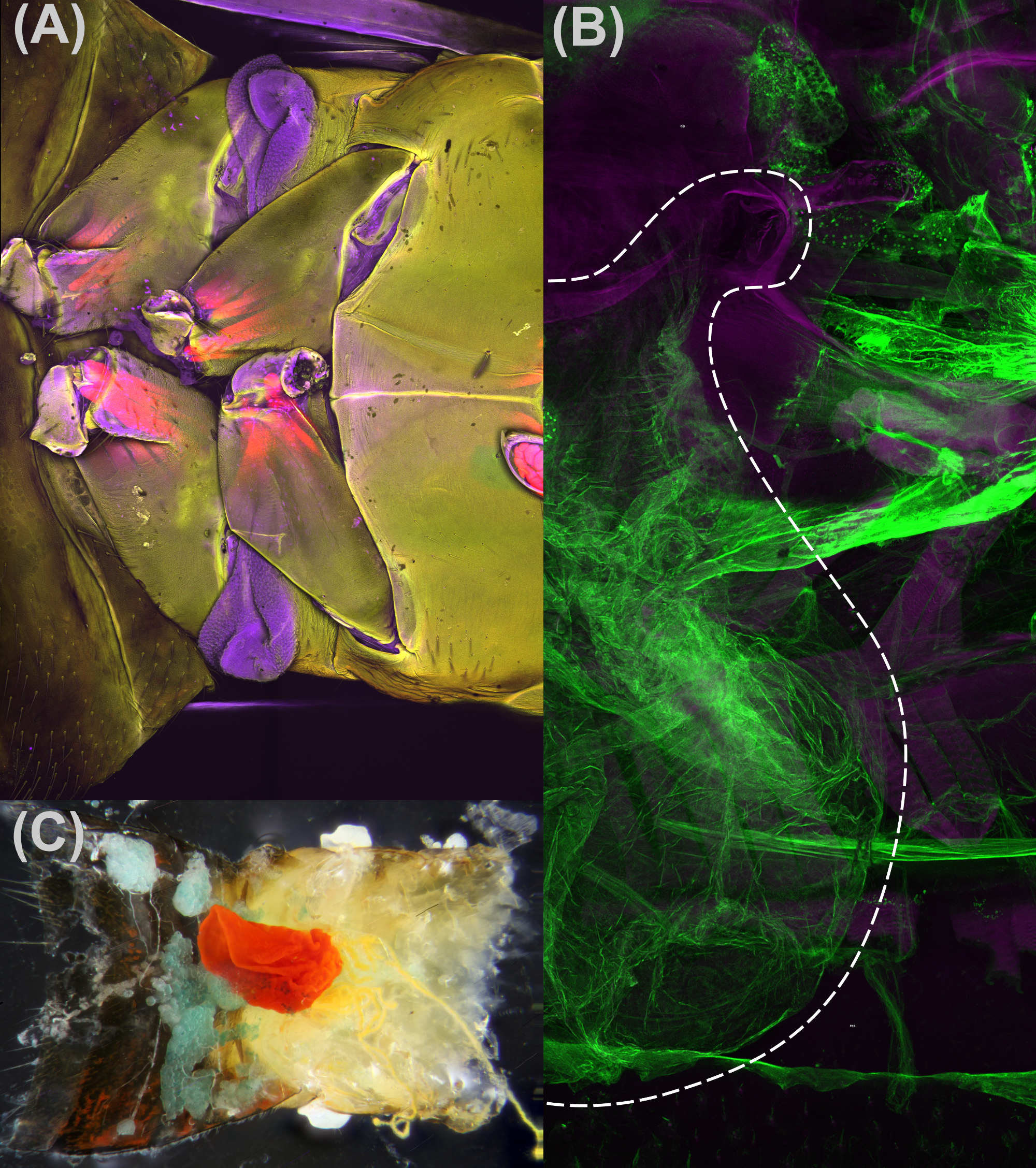
